# Working memory constraints for visuomotor retrieval strategies

**DOI:** 10.1101/2024.02.13.580155

**Authors:** Carlos A. Velázquez-Vargas, Jordan A. Taylor

## Abstract

Recent work has shown the fundamental role that cognitive strategies play in visuomotor adaptation. While algorithmic strategies, such as mental rotation, are flexible and generalizable, they are computationally demanding. To avoid this computational cost, people can instead rely on memory retrieval of previously successful visuomotor solutions. However, such a strategy is likely subject to strict stimulus-response associations and rely heavily on working memory. In a series of five experiments, we sought to estimate the constraints in terms of capacity and precision of working memory retrieval for visuomotor adaptation. This was accomplished by leveraging different variations of visuomotor item-recognition and visuomotor rotation recall tasks where we associated unique rotations with specific targets in the workspace and manipulated the set size (i.e., number of rotation-target associations). Notably, from Experiment 1 to 4, we found key signatures of working memory retrieval and not mental rotation. In particular, participants were less accurate and slower for larger set sizes and less recent items. Using a Bayesian-latent mixture model, we found that such decrease in performance is the result of both an increase in guessing behavior and of less precise samples from memory. In addition we estimated that participants’ working memory capacity was limited to 2-5 items, after which guessing increasingly dominated performance. Finally, in Experiment 5, we showed how the constraints observed across Experiments 1 to 4 can be overcome when relying on long-term memory retrieval. Our results point to the opportunity of studying other sources of memories where visuomotor solutions can be stored (e.g., episodic memories) to achieve successful adaptation.

## INTRODUCTION

Adapting motor output in response to unexpected sensory feedback or changing environmental demands is an essential process for skillful motor execution (Wolpert and Ghahramani 2000; Shadmehr et al., 2010; Krakauer et al., 2019). While this process of sensorimotor adaptation was originally thought to be the result of a low-order, implicit process (Jordan and Rumelhart, 1992; Squire and Zola, 1996; Squire, 2004; Mazzoni and Krakauer, 2006), in recent years it has become clear that higher-order cognitive strategies play a considerable role (Tsay et al., 2023; Huberdeau, et al., 2015; McDougle et al., 2016). In fact, implicit adaptation processes appear to be incapable of overcoming perturbations in a number of situations (Bond and Taylor, 2015; Morehead et al., 2017; Kim et al., 2019; Hadjiosif et al., 2021) and the use of cognitive strategies is necessary to improve performance (Wilterson and Taylor, 2021). Despite the mounting evidence for the importance of strategies in sensorimotor adaptation, we currently know very little about the underlying cognitive processes that support them.

In a prior study, we found evidence that people can employ (at least) two broad classes of strategies in a visuomotor rotation task: an algorithmic strategy that involves the mental simulation of different aiming solutions to overcome the rotation and a retrieval strategy that can “cache” previously successful aiming solutions (McDougle and Taylor, 2019). Algorithmic strategies, operationalized as a form of mental rotation (Pellizer and Georgopoulos, 1993), can be flexible and generalizable (McDougle and Taylor, 2019; Stransky et al., 2010), but come at a computational cost. In particular, reaction times (RTs) linearly increase with the rotation magnitude (Shepard and Metzler, 1971; McDougle and Taylor, 2019), indicative of the higher computational demands when the strategy is performed for longer – similar to the depth of tree search in planning. Indeed, people have a tendency to avoid situations that require greater mental rotation (Morsella et al., 2011).

Alternatively, participants could forgo the computational cost of an algorithmic strategy by attempting to retrieve a previously successful aiming solution from a short-term memory cache (Haith and Krakauer, 2018; Fresco et al., 2022). In effect, participants could construct a stimulus-response look-up table between a target location and its corresponding aiming solution. Compared with algorithmic strategies, this greatly reduces the time it takes to implement the strategy and reduces movement variance around the solution (McDougle and Taylor, 2019). However, retrieval strategies are likely subject to capacity constraints of short-term memory: Participants may be able to cache strategies when task complexity (set size) is low but this approach may break down as complexity increases, based on studies of working memory capacity outside the motor domain (Miller, 1956; Oberauer et al., 2016; Oberauer et al., 2018; Cowan, 2017).

Potential constraints of memory capacity for retrieval strategies could significantly limit their usefulness for visuomotor adaptation over the long term. Previous studies have shown that strategy implementation improves with training, in terms of speed and accuracy of execution (Huberdeau et al., 2019; McDougle and Taylor, 2019; Wilterson and Taylor, 2021). However, it remains an open question whether this is due to greater efficiency in the algorithmic computations for visuomotor mental rotation or whether it reflects a shift toward cached solutions being retrieved from memory (Logan, 1988). In classic Shepard-like mental rotation tasks, improvements in performance are thought to be the result of item-based memory retrieval for familiar stimuli rather than improvement in the algorithmic computations themselves (Kail, 1986; Tarr and Pinker, 1989; Heil et al, 1998). These findings are consistent with the Instance Theory of Automatization of skills where early in learning tasks are performed using computationally-demanding algorithmic processes but slowly give way to less demanding retrieval-based memory processes for familiar stimulus-response associations (Logan, 1988). Indeed, we have found that participants readily switch from an algorithmic strategy to a retrieval over the course of training; however, this was only observed for conditions involving a few training targets in the workspace, suggesting that memory capacity limitations may constrain strategy selection and, ultimately, their efficacy for visuomotor adaptation (McDougle and Taylor, 2019).

Here, in a series of studies, we sought to determine if retrieval strategies for visuomotor adaptation are subject to constraints of working memory, in terms of capacity and precision. While previous studies have addressed memory retrieval processes in visuomotor tasks (Pellizzer and Georgopoulos, 1993), and more recently in visuomotor rotation experiments (McDougle and Taylor, 2019), they have remained agnostic to situations where visuomotor solutions are experienced a single, or few, times. Such scenarios are a key component of human experience and have been shown to guide decision-making in a wide variety of settings (Yoo and Collins, 2021; Gershman and Daw, 2017; Bornstein et al., 2017).

To address this question, we made use of the visuomotor rotation task, which has not only served as a model paradigm for studying adaptation (Krakauer, 2009; Pine et al., 1996) but is also ideally suited to study cognitive strategies since the solution is expressible in the relevant dimension of the task (Taylor et al., 2014). We combine this adaptation task with classic experimental approaches, such as item-recognition (Sternberg 1966; Pellizzer and Georgopoulos, 1993), probed recall (Murdock, 1968), and computational models to study visuospatial working memory (Zhang and Luck, 2008; Adam et al., 2017; Lee and Wagenmakers 2014). Through this set of experiments, we find that retrieval strategies are constrained to a range of 2-5 items, consistent with the findings of visuospatial working memory, but can be overcome via long-term memory following substantial repetition. These findings suggest that retrieval strategies may be useful for visuomotor adaptation over the long-term.

## RESULTS

### Experiment 1

#### Behavioral results

The goal of our first experiment was to verify that people can leverage retrieval strategies, instead of an algorithmic strategy (i.e., mental rotation), to successfully solve a visuomotor rotation task and to estimate the precision of retrieval strategies. To test this, participants (n=15) performed a visuomotor item-recognition task (Sternberg, 1969; Pellizzer and Georgopoulos, 1993), where they first observed a sequence of targets, ranging from 2-5 targets (set size), that were displayed one at a time along a ring (Figure 1A; see *Materials and Methods* for details). Then, a cued target from the sequence was presented, and the goal was to reach to the location of the subsequent target in the sequence. Within the item-recognition task, we embedded a visuomotor rotation task by manipulating the angular rotation between the endpoint cursor feedback and the end of the reach. Importantly, the angular difference between the cued and subsequent target were varied to simulate a visuomotor rotation, which ranged from −90° to 90°. This way, if participants correctly reached to the subsequent target in the sequence, they would also hit the cued target with the cursor. We hypothesized that if participants were employing retrieval-based strategies, then RTs to the subsequent targets would not vary with the angular difference between the cued and subsequent targets (i.e., visuomotor rotation magnitude). After a given sequence was experienced, the relationship between the target locations and rotations was randomized for the next sequence.

**Figure 1:**
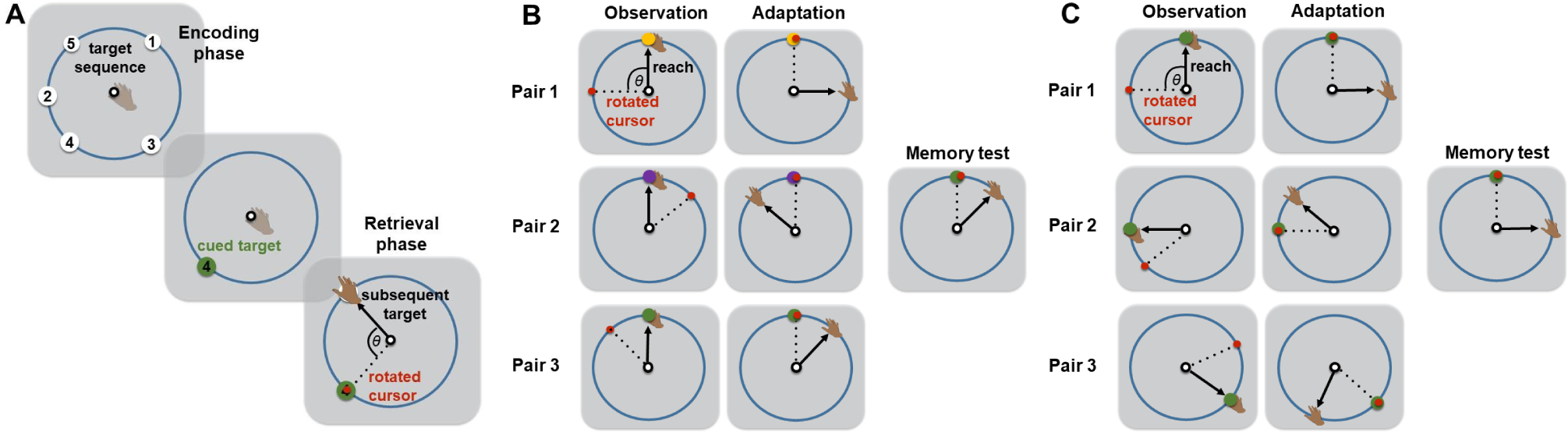
**(A)** Visuomotor item-recognition task from Experiment 1 and 2. A sequence of targets appeared, one at a time, on a circle followed by the presentation of one target from that sequence. The participants had to reach the location of the subsequent target on the sequence. **(B)** Visuomotor rotation recall task from Experiment 3. A sequence of pairs of trials was shown. In the first trial of each pair, the participants reached to the colored target and observed the rotation associated with it (observation). In the next trial, they attempted to counteract the rotation (adaptation). In the memory test, one colored target from the ones in the sequence was shown (target of pair 3 is shown as an example) and participants had to counteract the rotation associated with it. **(C)** Visuomotor rotation recall from Experiment 4. In this experiment each rotation value was associated with a different target location instead of colors as in Experiment 3. Target of pair 1 is shown as an example of the memory test.

Overall, participants successfully reached the subsequent target across the different rotation magnitudes as reflected in a significant positive correlation between the adapted magnitude and the rotation magnitude (r^Pearson^(178) = 0.96, p < 0.001; r^Spearman^(178) = 0.96, p < 0.001; Figure 2A). However, accuracy decreased with the sequence length as suggested by larger unsigned errors for longer sequences (r^Pearson^(58) = 0.74, p < 0.001; r^Pearson^(58) = 0.79, p < 0.001; Figure 2B), providing preliminary evidence that the precision of retrieval-based strategies are subject to set size constraints.

**Figure 2:**
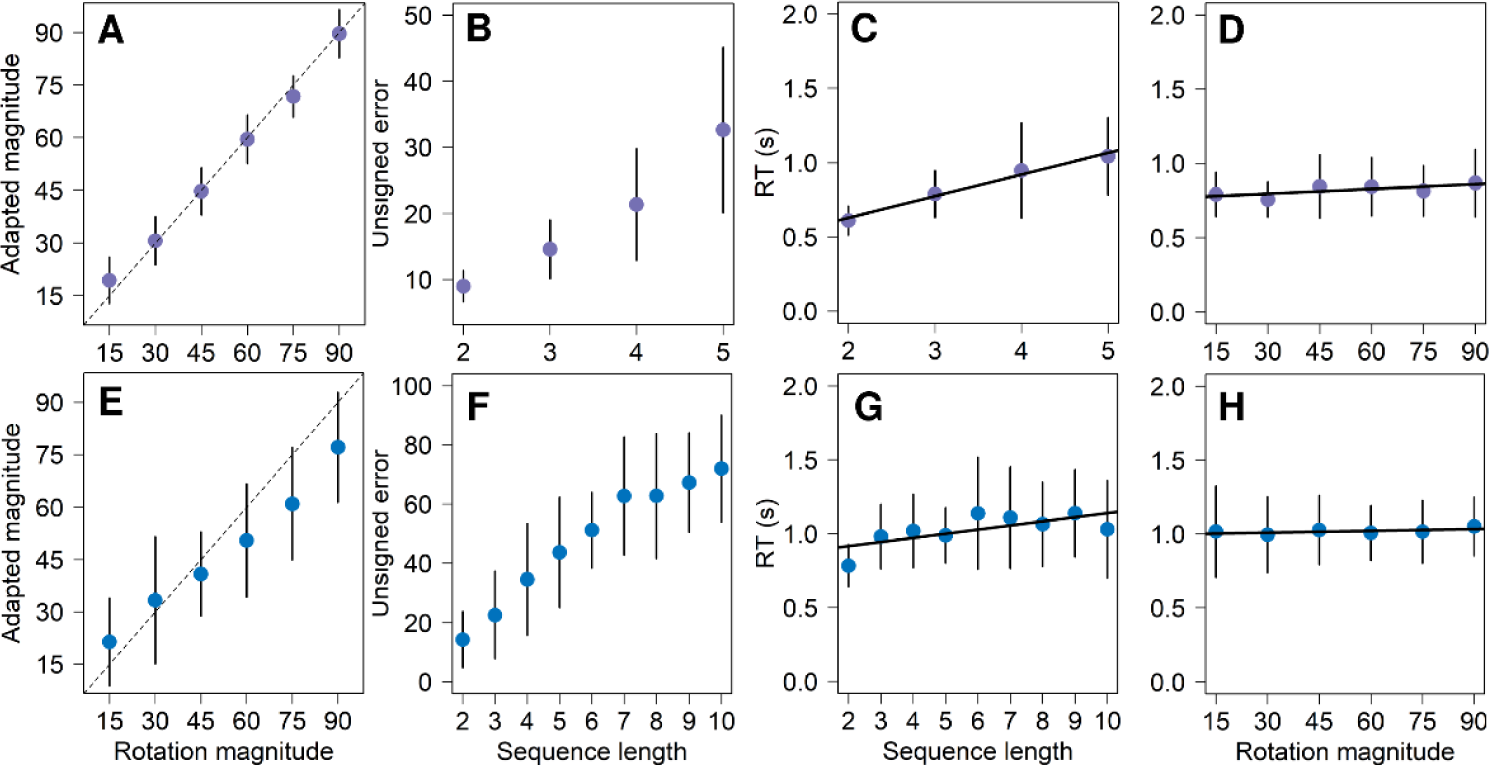
Behavioral results of Experiment 1 (purple) and Experiment 2 (blue). **(A)** and **(E)** Per-subject mean of adapted magnitude for each rotation magnitude. The dashed lines indicate performance when perfectly counteracting the rotation. **(B)** and **(F)** Per-subject mean of unsigned errors for each sequence length. **(C)** and **(G)** Per-subject median of reaction times for each sequence length. **(D)** and **(H)** Per-subject median of reaction times for each rotation magnitude. Black solid lines on RTs plots show the linear models fitted to the data. Error bars represent the standard deviation with respect to the mean.

A key prediction of this study was that RTs would increase with the sequence length (Pellizzer and Georgopoulos, 1993) but not the rotation magnitude (McDougle and Taylor, 2019) if the participants were performing memory retrieval instead of mental rotation. Indeed, we found that sequence length (*β* = 0.14, p < 0.001, R^2^ = 0.35, F(1, 58) = 32.18; Figure 2C), but not rotation magnitude (*β* = 0.001, p = 0.15, R^2^ = 0.02, F(1, 88) = 2.02; Figure 2D), significantly predicted the median of participants’ RTs in a linear regression analysis.

As a key property of working memory retrieval, we looked for serial position effects on performance and found that unsigned errors were indeed smaller (r^Pearson^ (58) = −0.57, p < 0.001; r^Spearman^(58) = −0.55, p < 0.001; Figure S1A) and RTs lower (r^Pearson^ (58) = −0.40, p = 0.001; r^Spearman^(58) = −0.50, p < 0.001; Figure S1E) for more recent targets in the sequence (i.e., a recency effect).

#### Modeling results

Following previous work on spatial working memory (Zhang and Luck, 2008; Adam et al, 2017), we analyzed the target error distributions in our task (Figure 3A) using a Bayesian latent-mixture model to better characterize the precision of retrieval strategies (Lee, 2018; Lee and Wagenmakers, 2013; Shiffrin et al, 2008; Figure 3C). In particular, we assumed that target errors were generated by either a memory distribution, represented by a Von Mises distribution, or a guessing distribution, represented by a Uniform distribution. The Von Mises distribution is similar to a Normal distribution but adapted to circular data given the structure of our reaching tasks. We implemented this model at the group level in order to have enough data points to generate reliable estimates of the distribution of the parameters. For our analyses, we focused on the group guessing rate parameter (θ) and the group memory precision parameter (κ) of the distribution (See *Materials and Methods* for details). The κ parameter is similar to the standard deviation in the Normal distribution, however, lower values of κ indicate higher dispersion of the data.

**Figure 3:**
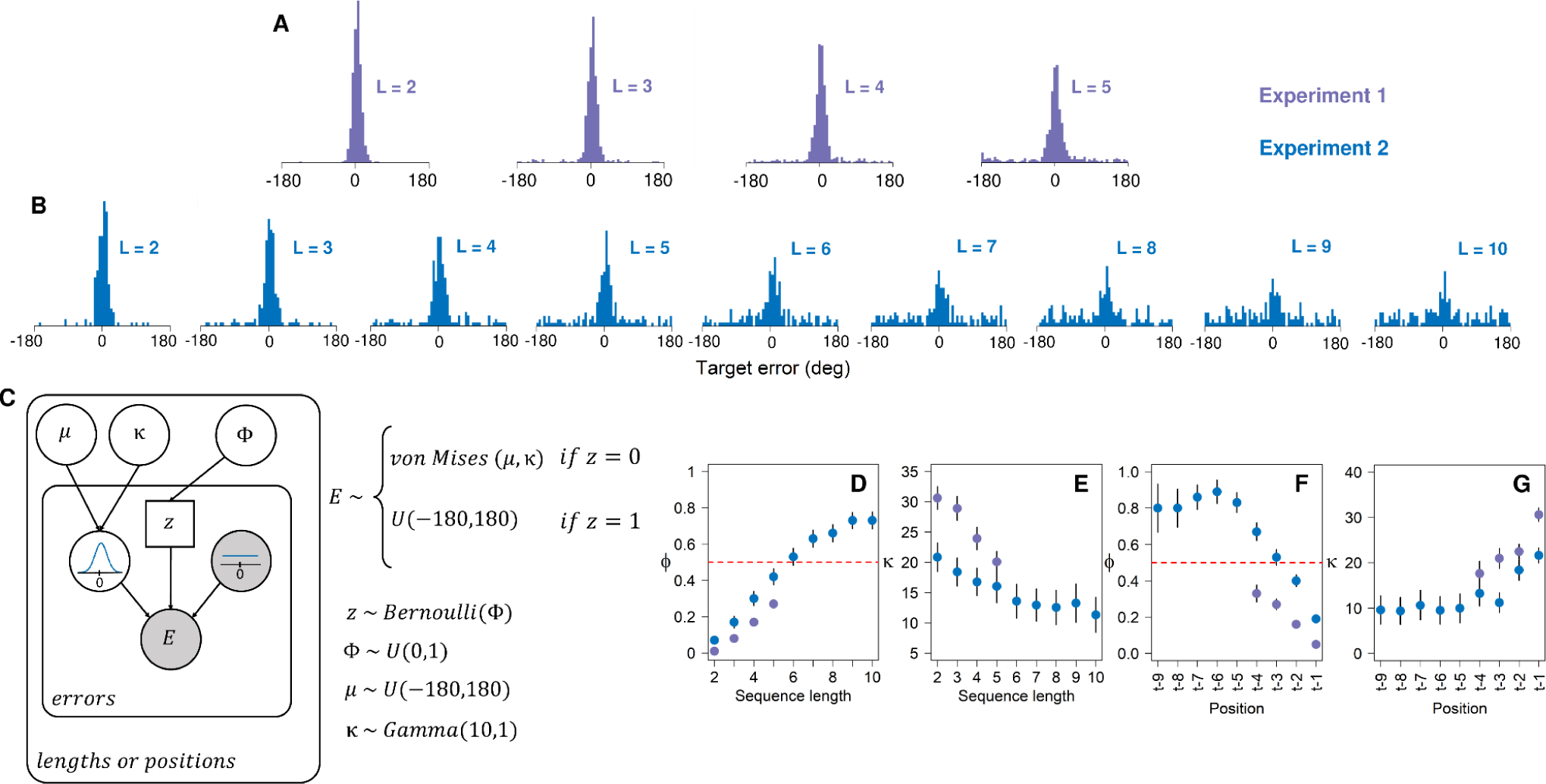
Analysis of target error distribution using a Bayesian latent-mixture model. **(A)** and **(B)** Pooled target error distributions for each sequence length of Experiment 1 and Experiment 2. **(C)** Graphical representation of the Bayesian latent-mixture model of target errors (See *Materials and Methods* for details). **(D)** Posterior distribution of the group guessing rate parameter, ɸ, for each sequence length. The red dashed line indicates ɸ = 0.5. **(M)** Posterior distribution of the group memory precision parameter, κ, for each sequence length. **(N)** Posterior distribution of the group guessing rate parameter, ɸ, for the last (t-1), second to last (t-2), and so on, targets across sequence lengths. Error bars represent the standard deviation with respect to the mean.

As expected from previous studies (Adam et al, 2017), we found that the guessing rate increased with the sequence length (r^Pearson^ = 0.98, 95%CI [0.97, 0.98]; r^Spearman^ = 0.96, 95%CI [0.96, 0.96]; Figure 3D), while the precision of the memory distribution decreased (r^Pearson^ = −0.89, 95%CI [−0.86, −0.92]; r^Spearman^ = −0.89, 95%CI [−0.86, −0.92]; Figure 3E). These modeling results are consistent with the decrease in accuracy observed in participants’ unsigned errors (Figure 2B) and the pooled target error distributions (Figure 3A). Furthermore, we were interested in whether the guessing rate and the precision of the memory distribution would vary depending on the position of the target in the sequence (i.e. serial position effects). Indeed we found a negative correlation between the target position in the sequence and the guessing rate θ (r^Pearson^ = −0.95, 95%CI [−0.94, −0.97]; r^Spearman^ = −0.94, 95%CI [−0.92, −0.96]; Figure 3F), indicating that across sequence lengths, participants guessed less for more recent targets. Furthermore, we found a positive correlation between the memory precision κ and the position of the target, such that there was higher precision for more recent items (r^Pearson^ = 0.87, 95%CI [0.82, 0.90]; r^Spearman^ = 0.85, 95%CI [0.79, 0.90]; Figure 3G).

Together, the findings of Experiment 1 provide evidence that people can use a retrieval strategy, and not mental rotation, to successfully solve a visuomotor rotation task. In addition, we showed that retrieval is subject to capacity limitations as reflected in larger errors, higher guessing rates, lower memory precision and higher reaction times for longer sequences. Furthermore, we found serial position effects – a signature of working memory retrieval – where people had smaller errors, were faster, guessed less and had more precise memories for more recent targets in the sequence. Notably, even for sequences of length five, guessing represented a relatively low proportion of the trials (posterior mean of θ for length five = 0.27), which suggests that the memory capacity of participants may extend beyond five targets. In Experiment 2, we explore this limit by exposing participants to longer sequences (larger set size).

### Experiment 2

#### Behavioral results

The goal of Experiment 2 was to explore the limits of visuomotor retrieval beyond the sequence lengths tested in Experiment 1. Specifically, we sought to replicate our findings for sequences up to five targets and to assess whether the trends in our relevant variables (unsigned errors, RTs, guessing rate and memory precision) would extend to sequences of up to 10 targets.

As in Experiment 1, we found a significant positive correlation between the adapted magnitude and the rotation magnitude (r^Pearson^ (154) = 0.76, p < 0.001; r^Spearman^ (154) = 0.78, p < 0.001; 2E), confirming that participants successfully reached the subsequent target in the sequence, and therefore counteracted the rotation. Additionally, we found that participants’ accuracy decreased when they faced more targets as reflected in larger unsigned errors for longer sequences (r^Pearson^(115) = 0.74, p < 0.001; r^Spearman^(115) = 0.75, p < 0.001; 2F). Interestingly, when we compared the unsigned errors between Experiment 1 and Experiment 2 over the same sequence lengths (two to five targets), we found that participants from Experiment 2 performed significantly worse. This difference was tested using 2X4 repeated measures ANOVA, having as factors the experiment number and sequence length, and revealing a significant main effect of the experiment number (F(1,104) = 16.31, p < 0.001, η^2^= 0.08).

Similar to Experiment 1, RTs linearly increased with sequence length (*β* = 0.02, p = 0.006, R^2^ = 0.06, F(1, 115) = 7.68; 2G) but not rotation magnitude (*β* = 0.006, p = 0.68, R^2^ = 0.002, F(1, 76) = 0.16; 2H), supporting a memory retrieval strategy instead of mental rotation. Additionally, we found that participants in Experiment 2 had significantly higher RTs than participants in Experiment 1 over the same sequence lengths (two to five targets). This difference was assessed using 2X4 repeated measures ANOVA, where experiment number and sequence length were the factors, and finding a significant main effect of the experiment number (F(1,104) = 5.41, p = 0.02, η^2^= 0.035).

Corroborating the serial position effects of Experiment 1, we found that participants had smaller unsigned errors (r^Pearson^ (112) = −0.56, p < 0.001; r^Spearman^(112) = −0.56, p < 0.001; Figure S1B) and lower RTs (r^Pearson^ (112) = −0.31, p < 0.001; r^Spearman^(112) = −0.28, p = 0.002; Figure S1F) for more recent targets in the sequence. We can observe a similar serial position effect in the target error distributions (Figure S2).

#### Modeling results

Following the logic of Experiment 1, we analyzed the source of the target errors (Figure 3B) using a Bayesian latent-mixture model. Again, we found that the guessing rate θ increased with the sequence length (r^Pearson^ = 0.96, 95%CI [0.95,0.96]; r^Spearman^ = 0.96, 95%CI [0.95,0.97]; Figure 3D), this time following a sigmoidal-like shape. Furthermore, we found that guessing became predominant (occurring >50% of the trials) for sequences above length five, which was estimated by fitting a sigmoidal function (f) to the posterior means of Figure 2L and later computing f^−1^(0.5), which estimated 5.6 targets (Figure S3A). Similarly, based on the function (f), we found that the asymptote of the guessing rate (θ) was at 0.74. Regarding memory precision (κ), we found a negative correlation with the sequence length (r^Pearson^ = −0.70, 95%CI [−0.61,-0.77]; r^Spearman^ = −0.70, [−0.62,-0.77]; Figure 3E), confirming the decay in memory precision with the sequence length of Experiment 1. In addition, by visual inspection, we corroborated the behavioral differences between Experiment 1 and Experiment 2, showing that participants from Experiment 2 performed worse over the same sequence lengths: they guessed more and had less precise memories in target sequences with two to five targets (Figure 3D - 3G).

Finally, we performed our mixture analysis to identify serial position effects, finding a negative correlation between the guessing rate θ and the position of the target in the sequence (r^Pearson^ = −0.81, 95%CI [−0.77,-0.85]; r^Spearman^ = −0.77, [−0.72,-0.82]; Figure 3F), indicating that people guess less for the most recent items across all sequences (recency effect) as in Experiment 1. Additionally, we found a positive correlation between memory precision κ and the sequence length (r^Pearson^ = 0.68, 95%CI [0.61,0.74]; r^Spearman^ = 0.64, 95%CI [0.57,0.71]; Figure 3G), suggesting that they had more precise memories for more recent items.

In summary, the results of Experiment 2 corroborated the main findings of Experiment 1 regarding the memory constraints of visuomotor retrieval strategies. Participants had larger errors, higher RTs, guessed more and had less precise memories for longer sequences. In addition, we confirmed a recency effect in unsigned errors, RTs, guessing rate, and the memory precision. As in Experiment 1, we found no evidence that the participants were performing mental rotation. Interestingly, however, participants in Experiment 2 performed significantly worse than participants in Experiment 1 over the same sequence lengths according to behavioral and model-based measures, which may be a result of generally higher task demands for longer sequences. Finally, we found that the asymptote of guessing was 74% of the trials, and that guessing began to dominate performance (occurring >50% of the trials) for sequences above 5 targets.

The goal of Experiments 1 and 2 was to embed a visuomotor rotation into the item-recognition task of Pellizer and Georgopoulos (1993) to verify the ability of retrieval strategies to solve a visuomotor rotation task. Admittedly, the design of Experiments 1 and 2 depart from what is typically required in a visuomotor rotation task. Nonetheless, they serve as a bridge to study working memory constraints for a retrieval strategy in a more standard visuomotor rotation task, which we systematically build toward in the following set of studies.

### Experiment 3

#### Behavioral results

The goal of this experiment was to estimate the working memory capacity and precision of a retrieval strategy in a task that is a step closer to a standard visuomotor rotation task. In this experiment, we implemented a trial-pair design (McDougle and Taylor, 2019; Figure 1B) where participants (n=15) were first asked to reach toward a single target location, always at 90°, and observe a cursor rotation. In the following trial, they were tasked with counteracting the rotation, therefore making the cursor hit the target. Participants were exposed to sequences of such trial-pairs with lengths ranging from one to five pairs (i.e., set size). Importantly, each pair was associated with a unique rotation which was indicated with a distinctive target color – a common dimension studied in spatial working memory. The rotations could take the values of ±30, ±60 or ±90. At the end of the sequence presentation, there was a memory test where one of the colored targets from the observed sequence was presented and participants had to counteract the rotation associated with it. Following each memory test, the association between the color of the targets and rotation magnitudes were changed (see *Materials* and *Methods* for details).

Similar to the visuomotor item-recognition studies, we found that in the memory test participants successfully counteracted the rotations as observed in a positive correlation between the adapted magnitude and the rotation magnitude (r^Pearson^(43) = 0.82, p < 0.001; r^Spearman^(43) = 0.83, p < 0.001; Figure 4A). However, participants had larger unsigned errors for longer sequences (r^Pearson^(73) = 0.64, p < 0.001; r^Spearman^(73) = 0.64, p < 0.001; Figure 4B). As in the earlier item-recognition tasks, we found that RTs linearly increased with the sequence length (*β* = 0.22, p < 0.001, R^2^ = 0.25, F(1, 73) = 24.5; Figure 4C) but not the rotation magnitude (*β* = −0.08, p = 0.43, R^2^ = 0.01, F(1, 43) = 0.61; Figure 4D), indicating that participants did not perform mental rotation but memory retrieval.

**Figure 4:**
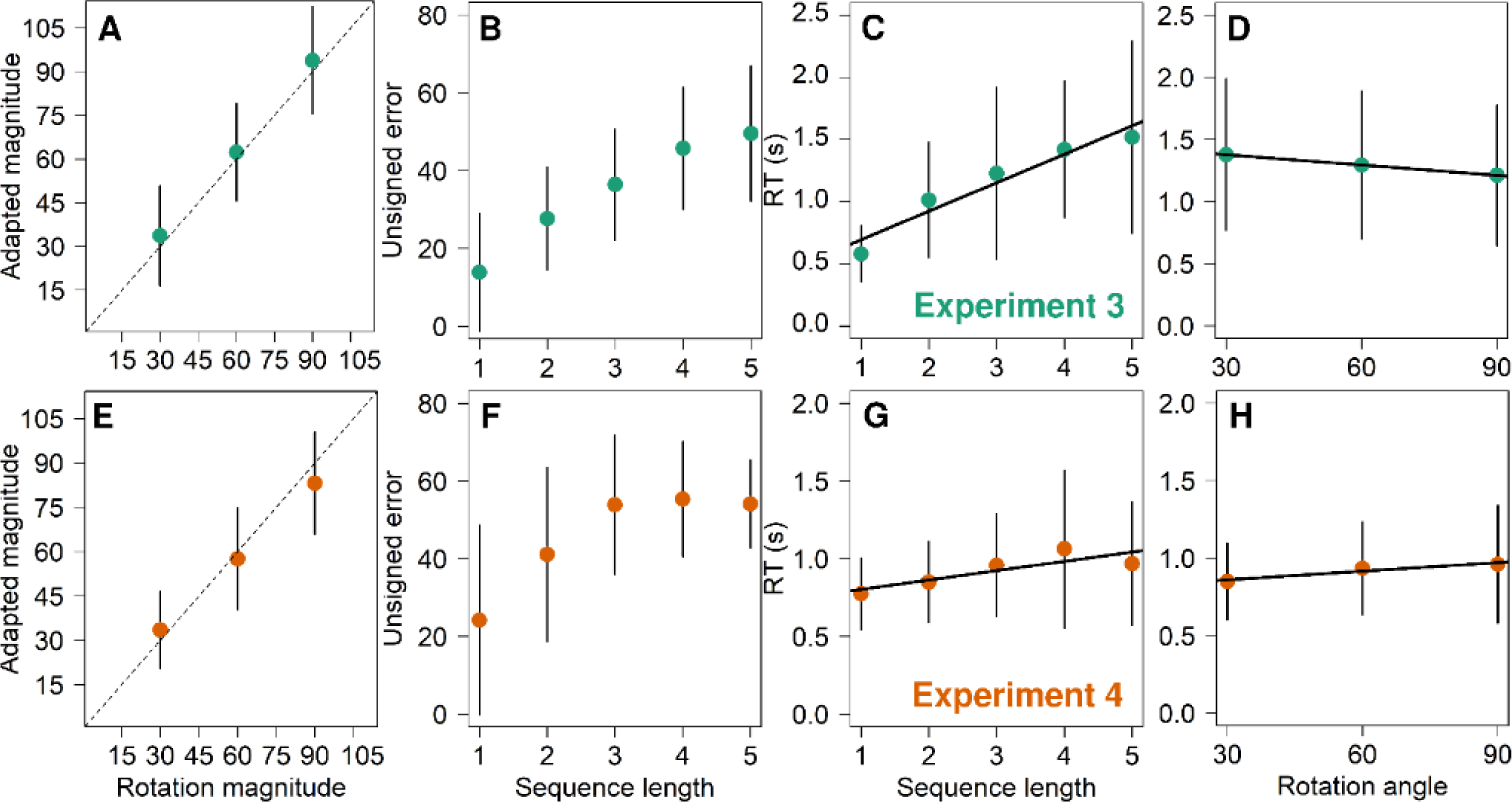
Behavioral results of Experiments 3 (green) and Experiment 4 (orange). **(A)** and **(E)** Per-subject mean of adapted magnitude for each rotation magnitude. The dashed lines indicate performance when perfectly counteracting the rotation. **(B)** and **(F)** Per-subject mean of unsigned errors for each sequence length. **(C)** and **(G)** Per-subject median of reaction times for each sequence length. **(D)** and **(H)** Per-subject median of reaction times for each rotation magnitude. Black solid lines show the linear models fitted to the data. Error bars represent the standard deviation with respect to the mean.

In addition, we found serial position effects where participants had smaller errors (r^Pearson^ (73) = −0.57, p < 0.001; r^Spearman^(73) = −0.56, p < 0.001; Figure S1C) and lower RTs (r^Pearson^ (73) = −0.25, p = 0.02; r^Spearman^(73) = −0.32, p = 0.004; Figure S1G) for more recent targets in the sequence.

#### Modeling results

We separated the source of the target errors (Figure 5A) using the same Bayesian latent-mixture analysis as in Experiment 1 and Experiment 2. Similarly, we found that the guessing rate θ increased for longer sequences (r^Pearson^ = 0.94, 95%CI [0.92,0.95]; r^Spearman^ = 0.95, [0.93,0.96]; Figure 5B) while the memory precision κ decreased (r^Pearson^ = −0.85, 95%CI [−0.80,-0.89]; r^Spearman^ = −0.88, [−0.84,-0.91]; Figure 5C). In addition, by fitting a linear model, f, to the posterior means of Figure 5B, we found that guessing began to dominate performance (>50% of trials) for sequences above 4 targets – specifically, f^−1^(0.5) = 4.16 (Figure S2B). Furthermore, we found serial position effects, where participants guessed less (r^Pearson^ = −0.85, 95%CI [−0.81,-0.88]; r^Spearman^ = −0.83, [−0.78,-0.88]; Figure 5D) and had more precise memories (r^Pearson^ = 0.78, 95%CI [0.73, 0.83]; r^Spearman^ = 0.76, [0.69, 0.83; Figure S4A) for more recent targets in the sequence.

**Figure 5:**
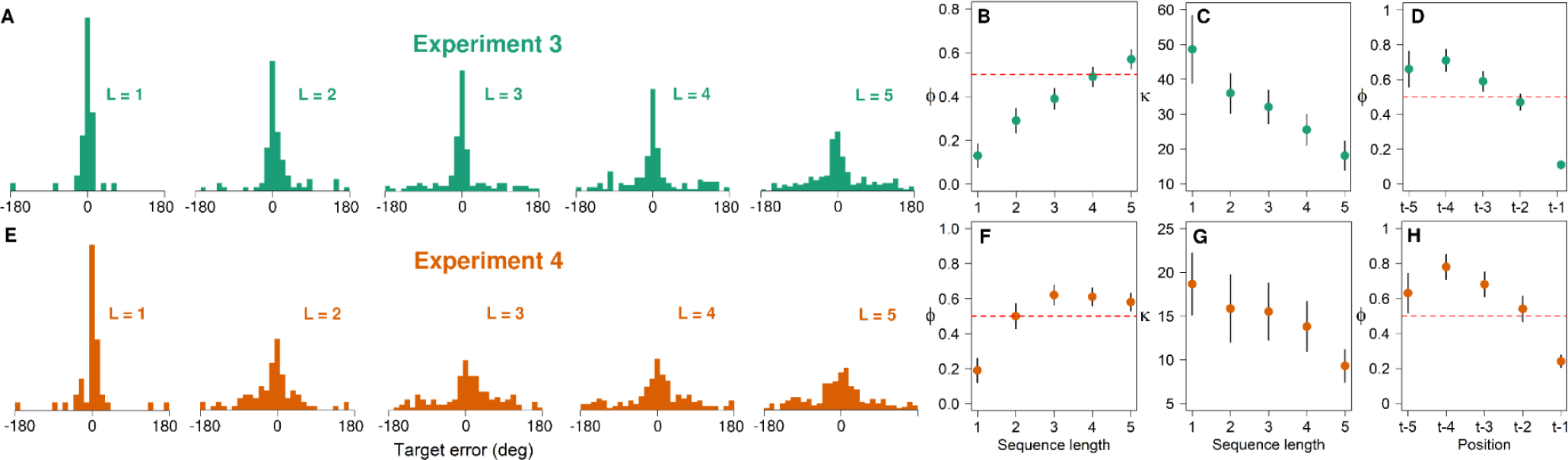
Modeling results of Experiment 3 (green) and Experiment 4 (orange) **(A)** and **(B)** Pooled target error distributions for each sequence length. **(B)** and **(F)** Posterior distribution of the group guessing rate parameter, ɸ, for each sequence length. The red dashed line indicates ɸ = 0.5. **(C)** and **(G)** Posterior distribution of the group memory precision parameter, κ, for each sequence length. **(D)** and **(H)** Posterior distribution of the group guessing rate, ɸ, for the last (t-1), second to last (t-2), and so on, targets across sequence lengths. Error bars represent the standard deviation with respect to the mean.

Overall, the results from Experiment 3 corroborate the main results in the visuomotor item-recognition tasks from Experiment 1 and Experiment 2, but in a task design that is a step closer to a standard visuomotor rotation task. In sum, we found that memory retrieval is constrained by the capacity of working memory as shown in larger unsigned errors, higher RTs, more guessing and lower memory precision for longer sequences. As in the visuomotor item-recognition task, we found no evidence that participants were performing mental rotation. Furthermore, we observed serial position effects both in our behavioral and model-based measures. Notably, in this experiment guessing becomes predominant for sequence lengths above 4 targets, a capacity similar to the item-recognition task of Experiment 2.

In the following studies we focused on memory retrieval associated with the target location instead of target color, a design that is more common in visuomotor rotation tasks.

### Experiment 4

#### Behavioral results

The goal of this experiment was to provide further evidence of the constraints of working memory retrieval in visuomotor rotation tasks; this time, by making participants associate rotation magnitudes with distinct target locations instead of colors. Associating rotation magnitudes with distinct target location has been used before on dual adaptation experiments (Baldeo and Henriques, 2013) as well as in trial-pair tasks (McDougle and Taylor, 2019). The target locations for every trial were randomly sampled without replacement out of eight possible values ranging from 0 to 315° in steps of 45. We used the same rotation magnitudes as in Experiment 3. Apart from this change, everything else remained as in Experiment 3 (see *Methods and Materials* for details).

As expected from our previous experiments, participants were able to successfully counteract the rotation indicated by a significantly positive correlation between the adapted magnitude and the rotation magnitude (r^Pearson^(43) = 0.79, p < 0.001; r^Spearman^(43) = 0.81, p < 0.001; Figure 4E). In addition, they had larger unsigned errors for longer sequence lengths (r^Pearson^(73) = 0.48, p < 0.001; r^Spearman^(73) = 0.44, p < 0.001; Figure 4F). Interestingly, participants in this experiment performed significantly worse than participants in Experiment 3 (i.e., color cues) as indicated by a 2X5 repeated measures ANOVA over unsigned errors, where experiment number and sequence length were the factors, and finding a significant main effect of the experiment number (F(1,140) = 15.9, p < 0.001, η^2^= 0.06). In addition, we found that RTs linearly increased with the sequence length (*β* = 0.06, p = 0.04, R^2^ = 0.05, F(1, 73) = 4.28; Figure 4G) but not rotation magnitude (*β* = 0.05, p = 0.33, R^2^ = 0.02, F(1, 43) = 0.95; Figure 4H), supporting memory retrieval over mental rotation.

Lastly, we also found serial position effects where participants had smaller errors (r^Pearson^ (73) = −0.39, p < 0.001; r^Spearman^(73) = −0.39, p < 0.001; Figure S1D) and marginally lower RTs (r^Pearson^ (73) = −0.22, p = 0.05; r^Spearman^(73) = −0.20, p = 0.07; Figure S1H) for more recent targets in the sequence.

#### Modeling results

When performing the Bayesian latent-mixture analysis to the target error distributions (Figure 5E), we found that the guessing rate was higher (r^Pearson^ = 0.73, 95%CI [0.69,0.77]; r^Spearman^ = 0.64, [0.56,0.71]; Figure 5F), while the memory precision was lower for longer sequences (r^Pearson^ = −0.66, 95%CI [−0.57,-0.75]; r^Spearman^ = −0.67, [−0.58,-0.76]; Figure 5G), both results are consistent with our previous experiments. In order to know when guessing behavior began to dominate participants’ performance (occurring >50% of the trials), we fitted a polynomial function, f, to the posterior means of Figure 5F. Supporting the behavioral differences between Experiment 3 and Experiment 4, we found that the guessing rate was greater than 50% of the trials starting from sequences of length two (f^−1^(0.5) = 1.98; Figure S3C), a noticeable reduction in capacity compared to Experiment 3, where this value was around four targets.

As in our previous experiments, there was a recency effect in the guessing rate θ and memory precision κ, where participants guessed less (r^Pearson^ = −0.71, 95%CI [−0.64,-0.78]; r^Spearman^ = −0.66, [−0.58,-0.74]; Figure 5H) and had, weakly, more precise memories (r^Pearson^ = 0.21, 95%CI [0.08, 0.33]; r^Spearman^ = 0.15, [0.05, 0.24]; Figure S4B) for more recent items, although primarily for the most recent item.

Overall, in Experiment 4 we were able to corroborate the main findings from our previous studies where participants’ performance decreased for longer sequences. However, compared to Experiment 3, where people associated rotation magnitudes to colors, in this experiment participants performed significantly worse, displaying a limited working memory capacity where guessing dominated performance starting from sequences of two targets. This finding would place a relatively low upper bound on how useful retrieval strategies can be for counteracting a visuomotor perturbation. Most visuomotor adaptation studies require participants to train at more target locations; however, the relationship between the target location and the rotation typically do not change from trial-to-trial, as in our trial-pair task. As such, participants experience many repetitions of the same stimulus-response association over the course of training, which would provide ample opportunity to store these associations for later retrieval according to the Instance Theory of Automatization (Logan 1988). In our final study, we addressed if these severe limitations in working memory capacity can be overcome, namely, by relying on a different memory storage: long-term memory.

### Experiment 5

#### Behavioral results

The goal of this final experiment was to test whether the retrieval strategies can overcome the limitations of working memory through repetition and, as such, long-term memory. To test this, participants (n=15) performed the trial-pair task as in Experiment 4, however, here they were exposed to sequences of five pairs, which would exceed our estimates of working memory capacity. Importantly, in contrast with Experiment 4, the rotation magnitudes associated with the target locations remained the same throughout the experiment but the order of the rotation-target pairs was randomized. This design allowed participants to experience the rotation-target associations multiple times, therefore, increasing the likelihood that they would be stored in long-term memory.

As in our previous experiments, we found that participants’ performance remained highly accurate across rotation magnitudes (r^Pearson^(43) = 0.96, p < 0.001; r^Spearman^(43) = 0.94, p < 0.001; Figure 6A). More interestingly, we observed that unsigned errors (r^Pearson^(103) = −0.39, p < 0.001; r^Spearman^(103) = −0.47, p < 0.001; Figure 6B) and RTs (r^Pearson^(103) = −0.33, p < 0.001; r^Spearman^(103) = −0.26, p = 0.007; Figure S5A) decreased over time, indicating that their performance gradually improved with training. Lastly, we found that RTs did not change with the rotation magnitude (*β* = 0.06, p = 0.53, R^2^ = 0.008, F(1, 43) = 0.38; Figure S5B) as indicated by a linear regression analysis, ruling out mental rotation.

**Figure 6:**
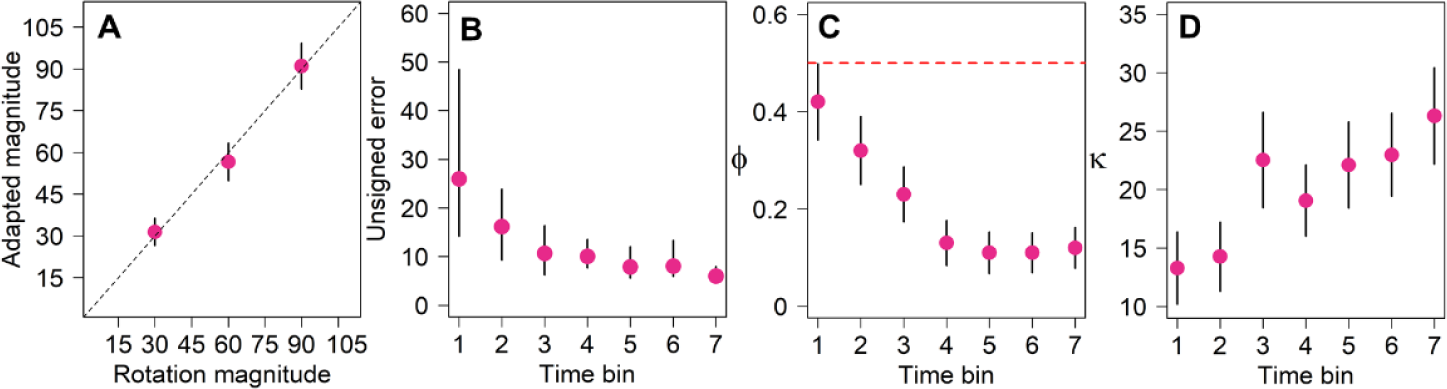
Behavioral and modeling results of Experiments 5. **(A)** Per-subject mean of adapted magnitude for each rotation magnitude. The dashed lines indicate performance when perfectly counteracting the rotation. **(B)** Per-subject mean of unsigned errors over time-bins. **(C)** Posterior distribution of the group guessing rate parameter, ɸ, over time bins. The red dashed line indicates ɸ =0.5. **(D)** Posterior distribution of the group memory precision parameter, κ, over time bins. Error bars represent the standard deviation with respect to the mean, except for (B) where error bars represent the interquartile range around the median due to non-normality of the data.

#### Modeling results

Corroborating our behavioral findings, we found that the guessing rate θ decreased over time (r^Pearson^ = −0.81, 95%CI [−0.78,-0.84]; r^Spearman^ = −0.79, [−0.74,-0.84]; Figure 6C) reaching a posterior mean of 12% of guessing by the end of the experiment. Notably, this value is lower than the guessing rate for sequences of a single target (i.e., the smallest set size) in Experiment 4 (posterior mean of θ for sequence length 1 = 19%). Similarly, the memory precision κ increased over time (r^Pearson^ = 0.70, 95%CI [0.62,0.77]; r^Spearman^ = 0.71, [0.63,0.77]; Figure 6D), reaching a posterior mean of κ = 26 by the end of the experiment, which is higher than the precision of target sequences of length 1 in Experiment 4 (posterior mean of κ for sequence length 1 = 13).

In summary, the results of Experiment 5 showed that the capacity limitations shown in Experiment 4, can be overcome over time, at a point where performance is better than in the shortest sequence relying on working memory.

### Model comparison

In order to validate the results from the Bayesian latent-mixture model, we compared its performance with a model that assumed that target errors had a single source, namely, the memory (Von Mises) distribution. We found that qualitatively and quantitatively –using leave-one-out cross validation (Vethari et al., 2017; Vethari et al., 2023) – the Bayesian latent-mixture model outperforms the single-source model in all experiments and sequence lengths (Figure S6).

## Discussion

### Summary

Through a series of studies we showed that participants can effectively use retrieval from memory as a strategy for visuomotor adaptation. From Experiment 1 to Experiment 4, we found key signatures of working memory retrieval, namely, lower accuracy and higher RTs as the number of targets in the training set increases (set size; Oberauer et al., 2018; Fukuda et al., 2010; Luck and Voguel, 1997; Sternberg, 1966, 1969). In addition, we found a recency effect where participants were more accurate and had lower RTs for more recent items, also a seminal finding in working memory studies (Glanzer and Cunitz, 1966; Oberauer et al., 2018). These behavioral results were supported by group model-based metrics, which revealed that participants were guessing more and had less precise memories for longer sequences (i.e., larger set size), as well as less guessing and more precise memories for more recent items – a recency effect. Experiment 5, revealed that these apparent working memory limitations could be overcome with repetition, suggesting that retrieval strategies can also rely on long-term memory. This later finding is consistent with the Instance Based Theory of Automatization and may provide clues as to how a strategy could lead to skill development (Logan, 1988).

### Capacity Limitations of Retrieval Strategies

From our model-based analysis, we found that guessing behavior began to dominate participants’ performance (more than 50% of trials) in the visuomotor item-recognition task for training sets beyond five targets, and similarly for the visuomotor rotation recall task with color cues, for sets beyond four targets. Both of these performance thresholds are similar to the capacity limitations documented in previous studies (Cowan, 2001, 2010). Interestingly, we found that for Experiment 4, where visuomotor rotation recall was based on location cues, guessing exceeded 50% of trials starting with sequences of two targets, which was a striking reduction in performance. This difference could have arisen given that, although spatial location and colors are both continuous variables, colors are usually associated with well known categories (red, green, purple, etc.), requiring little or no learning about them. On the other hand, target locations were not explicitly linked with well known categories in space, therefore, participants probably had to memorize the target locations as well as the rotation magnitude associated with them. Future experiments could test this hypothesis, for example, by providing clock “marks”, around the ring, indicating the locations where the target could appear. This way, participants can use a well known reference frame to remember the target locations (e.g. target at 6 linked with 90° rotation). We would expect that in this scenario performance would be similar to Experiment 3.

In addition, it is worth noticing that we found significant differences in performance between the two studies in the visuomotor item-recognition task. Specifically, in Experiment 2, participants showed overall lower performance, both in behavioral and model-based metrics, as compared to participants in Experiment 1 over the same set sizes. Since participants in Experiment 2 were additionally presented with sets greater than five and up to ten targets, we credit this effect to a higher cognitive load imposed by the average set size in the experiment. This decrease in performance has been reported before in working memory studies, although usually when participants’ perform a distracting task either from the same or a different domain (Vergauwe, 2010).

It has been widely debated whether human working memory is better described by models that assume a fixed-slot or a continuous resource capacity (Adam, Vogel and Awh, 2017; Ma, Husain and Bays, 2014). Whereas the original form of the fixed-slot models would assume a shift from retrieval to guessing after memory capacity is surpassed, the continuous resource models predict, instead, a decline in the memory precision as the number items to be remembered increases. Using the Bayesian latent-mixture model, we allowed for the incorporation of both of these assumptions. In particular, target errors were modeled as a combination of memory (Von Mises) distribution and a guessing (Uniform) distribution; at the same time, we allow the memory precision to be variable across set sizes. Interestingly, we found qualitative and quantitative evidence (Figure S5) that the decrease in participants’ performance was the result of a combination of both increased guessing and decreased memory precision, which is consistent with both frameworks. However, we did not find abrupt changes on participants’ performance, particularly on Experiment 2, where target sequences far exceeded typical capacity thresholds, which would be predicted by an all-or-none, fixed-slot model (Zhang and Luck, 2008). Instead, unsigned errors, guessing rate and memory precision, seem to describe continuous functions, results which better align with the predictions of continuous resource models (Wilken and Ma, 2004; Ma, Husain and Bays, 2014). Further work would be necessary to assess whether other instances of continuous resource models, e.g. that assume a different precision per trial (Ma, Husain and Bays, 2014), can provide a better description of this data set without the assumption of a guessing distribution.

Finally, we should note that we view these working memory capacity limitations as arising from limits in visuospatial working memory, given the remarkable similarity in our findings with studies of visual short-term memory in domains outside of motor control (Miller, 1956; Oberauer et al., 2016; Oberauer et al., 2018; Cowan, 2017; Adam, Vogel, and Awh 2017). This perhaps is not surprising since the reaching movements in our task were ballistic and visual feedback of the cursor was delayed to prevent implicit recalibration. While it is possible that proprioception or somatosensory information could be leveraged to recall a successful reach location with a specific target location (Sidarta et al., 2018; Kumar et al., 2021), the most salient features of the task were visuospatial – target and planned aiming location. Retrieval strategies could differentially rely on visuospatial and somatosensory “motor working memory”, which appear to be dissociable (Hillman and McDougle, 2024), depending on the nature of the task.

### Retrieval versus algorithmic strategies

In recent years, experimental work on visuomotor adaptation has revealed that people can deploy at least two kinds of cognitive strategies in response to feedback perturbations. Algorithmic strategies, which allow for the discovery of generalizable aiming solutions, are computationally demanding. On the other hand, memory retrieval allows for a fast and computationally effective way to recover known aiming solutions, but is limited in generalization and capacity. Previous work has shown that participants can switch from algorithmic to retrieval strategies as the experimental session progresses, which is reflected in a decrease in RTs over time (McDougle and Taylor, 2019). This transition is consistent with a process of automatization and memory consolidation characteristic of the development of a skill (Logan, 1988).

While retrieval strategies can convey an advantage in computational cost reflected in a reduction in RTs, this benefit is likely to become more evident as the aiming solutions are consolidated into long-term memory due to repetition. Indeed, in McDougle and Taylor (2019), RTs differences between participants performing algorithmic and retrieval strategies are accentuated as the aiming solutions are repeated over the experimental session. This enhancement in computational efficiency is analogous to the one of mental arithmetic (Fresco et al., 2023), where the solutions for common operations are readily retrieved from memory (e.g., the result of 5 times 5), whereas the solutions for less common, or novel, operations are more slowly performed algorithmically (e.g., the result of 23 times 11). We observed a similar improvement in computational efficiency in Experiment 5, where RTs, unsigned errors and guessing decreased, while memory precision increased, with more repetitions of the aiming solutions. However, this improvement was most likely driven by a transition from working memory to long-term retrieval and not from algorithmic computations to long-term retrieval, as we found no evidence of mental rotation in the experiment (Figure S5B).

However, when the aiming solutions are presented a single time as in Experiments 1-4, the computational efficiency gained through repetition is not attainable. In this scenario, where retrieval relies instead on working memory, the computational cost can increase as more solutions have to be stored. Evidence for this idea is found in previous work (Sternberg, 1966, 1969; Pellizzer and Georgopoulos, 1993) as well as in the present studies, where RTs linearly increase with the set size, which has been proposed to be the consequence of a scanning process over the items in memory (Sternberg, 1966, 1969; Donkin and Nosofsky, 2012). Notably, this linear increase in RT with set size in memory-based retrieval tasks mirrors the linear increase in RT with rotation magnitude in mental rotation tasks. However, there is substantial evidence suggesting that they are the result of fundamentally different psychological and neural operations (Pellizzer and Georgopoulos, 1993; McDougle and Taylor 2019; McDougle, Tsay, et al., 2023).

Whether memory scanning is computationally cheaper than mental rotation is an open question. From the set of studies we have presented, Experiment 4 has the most comparable design to McDougle and Taylor (2019) trial-pair rotation task, where the linear relation between RTs and rotation magnitudes was documented. Specifically, both designs vary the target location on which the aiming solution is tested. In our Experiment 4, participants reached RTs medians of around 1s for the longest target sequences, which is around the RTs values for the smallest rotations in McDougle and Taylor (2019), and where the largest rotations reached values of around 1.3s. This would suggest that retrieval, even when it occurs from working memory, could be computationally cheaper than mental rotation. However an experiment that evaluated these comparable designs could provide empirical evidence of the computational efficiency of each strategy.

### Automatization of a skill

It is well known that practice generally leads to task improvements in terms of speed and accuracy (Fitts and Posner, 1967). One potential explanation for this improvement is the increased efficiency of algorithmic processes (e.g. that mental rotation is performed faster and more accurately over time). Crucially, to the authors’ knowledge, there is currently not enough evidence that algorithmic processes, and in particular mental rotation, improve with practice (Tarr and Pinker 1989; Heil et al., 1998). Indeed, in McDougle and Taylor (2019) when participants were tasked with counteracting twelve different rotations associated with different targets, which favors the use of mental rotation over memory retrieval due to capacity limitations, their RTs only gently decreased with training, suggesting that participants continued using mental rotation with the same efficiency throughout the task.

On the other hand, it has been proposed that, when developing a new skill, people start by relying on algorithmic strategies but subsequently transition to perform memory retrieval of already known solutions (Logan, 1988) – with this transition entailing a reduction in the computational cost. While this process likely underlies a wide variety of human skills like mental arithmetic, we found evidence that long-term retrieval does not need to be preceded by algorithmic performance. Instead, it can result from consolidating task solutions that had already been stored in working memory. We can think, for example, that in a pool game, a player can temporarily store the shooting locations that were verbally or visually conveyed by a more experienced player without having to compute them themselves. These solutions, if successful, can be consolidated for future use into long-term storage. Such a strategy can prove successful in the short-term, only relying on algorithmic performance in the absence of temporarily stored solutions, such as in the presence of novel stimuli (Frank and Macnamara, 2019)

However, when the task at hand has no explicit incentive to use retrieval right from the beginning (as in our studies), a more natural transition in the development of a skill could be starting with algorithmic processes, followed by working memory retrieval and long-term retrieval if the solutions are consolidated (Atkinson and Shiffrin, 1968). The inability to transition from algorithmic to retrieval strategies when working memory capacity is exceeded (McDougle and Taylor, 2019), highlight the relevance for the study of the latter to understand the successful acquisition of visuomotor skills.

While more research is necessary to understand how memory retrieval contributes to visuomotor adaptation, and overall in motor learning, it is conceivable that such memories could also work as a starting point over which subsequent, implicit processes could operate. In effect, visuomotor solutions could come from verbal instructions or recent learning episodes, narrowing the solution space for implicit recalibration and speeding up learning. This work opens future avenues of research where other memories, such as episodic memories (Gershman and Daw, 2017; Bornstein et al., 2017; Tulving, 2002), can be the subject of study as potential sources of visuomotor solutions for adaptation.

## Materials and Methods

### Participants

73 undergraduate students (34 males, 37 females, 1 non-binary and 1 preferred not to say; mean age = 20, sd = 1.2) from Princeton University were recruited through the Psychology Subject Pool. Sample sizes were based on previous studies on visuomotor adaptation where the number of subjects per condition typically ranges from 10 to 20 participants per condition. The experiments were approved by the Institutional Review Board (IRB) and all participants provided informed consent before performing the experiment.

### Apparatus and task design

Participants performed horizontal, center-out movements holding a digital pen over a Wacom tablet. The movements were recorded at a sampling rate of 60 Hz in a 43.18cm, 1024X768, LCD Dell monitor running on Windows 7. Visual feedback of the hand was occluded by the monitor, which was mounted 25 cm above the tablet. On every trial, the participants attempted to find the monitor’s center (d = 5 mm). They were aided by a white ring that either expanded or contracted with the radial distance of the participant’s hand from the center. When the hand was 6 mm from the center, a red circular cursor (d = 4mm) appeared at the current hand position – the start position. After holding the start position for 1 s, a circular target (d = 7mm) appeared along a blue ring (d = 70mm). The participants were instructed to make ballistic center-out movements as fast and accurately as possible such as to slice through the blue ring, with the ultimate goal of hitting the target with the cursor. In order to achieve the latter, each experiment had varying requirements which will be described below. After leaving the start position, visual feedback of the cursor was removed. If the movement duration between leaving the start position and crossing the blue ring exceeded 0.6 s, the participants received an auditory warning (“too slow”). When the participants’ hand exceeded the blue ring radius, the cursor would show up along the ring. If it appeared on top of the target – a hit – the participants would hear a pleasant sound (“ding”); otherwise, they would hear an unpleasant sound (“buzz”) – a miss. The cursor feedback was delayed 0.5 s to reduce the possible role of implicit adaptation (Brudner et al., 2016) in the experiments. The cursor endpoint feedback remained on the screen for 0.5 s, after which participants would attempt to find the center to begin the next trial. Every experiment consisted of a series of blocks separated by a pause, where the message *“Wait for experimenter’s instructions”* would appear and participants would be told what to do in the next block. All experiments were adapted such that they did not exceed one hour.

### Experiment 1: Visuomotor item-recognition

In this experiment (n=15), we sought to establish that participants could use a retrieval strategy to solve a visuomotor rotation without using a visuomotor mental rotation strategy. Here, we embedded a visuomotor rotation task in the context of a “classic” working-memory item-recognition task (Sternberg 1969; Pellizzer and Georgopoulos 1993). The experiment unfolded over a series of encoding and retrieval phases. In the encoding phase, the participants first observed a sequence of white targets displayed one at a time along the blue ring, and were asked to memorize them (Figure 1A). Each target remained on the screen for 0.8s and was separated from the next one by 0.15 s. The angular location of the targets over the ring was randomly sampled with replacement out of 24 possible locations ranging from 0 to 345° in steps of 15°. After 0.2 s of the sequence presentation, the retrieval phase began. In this phase, one target was selected randomly from the sequence and displayed in green to the participants (“cued’’ target). The participants were instructed to perform a reaching movement to the location of the target that appeared immediately after it in the sequence (“subsequent” target). We manipulated two independent variables: the sequence length, which ranged from 2 to 5 targets, and the angular separation between the cued and subsequent target, which ranged from −90 to 90° in steps of 15° and excluding 0°, giving a total of 12 values. Importantly, the cursor endpoint position was rotated in the opposite direction and with the same magnitude as the angular separation between the cued and subsequent target (Fig. 1A). Therefore, if the participants correctly reached the position of the subsequent target, they would hit the cued target with the cursor – in effect, a visuomotor rotation task.

A crucial prediction from this experiment was that if participants were performing memory retrieval, their RTs would increase with the sequence length, indicative of memory scanning (Sternberg, 1969), but not with the rotation magnitude, indicative of mental rotation (McDougle and Taylor, 2019).

In order to make sure the participants understood the experiment, they underwent a series of preparation blocks where they first learned to find the center of the tablet and make straight reaching movements to the targets (48 trials); then, they were exposed to target sequences with all lengths and rotation magnitudes to get familiarity with the task (48 trials). Then, in the actual experiment, each sequence length and rotation magnitude combination was tested three times (144 trials). At the end, participants underwent a washout block (24 trials) where they were instructed to reach directly to targets while not receiving endpoint feedback. The experiment had a total of 264 trials.

### Experiment 2: Extended visuomotor item-recognition

Since guessing behavior in Experiment 1 consistently remained below 50% across all sequence lengths, we hypothesized that participants’ working-memory capacity exceeded the demands imposed by the task. Therefore, in Experiment 2 (n=13), we extended the design of Experiment 1 to include sequences up to length ten. We were particularly interested in whether the guessing behavior and memory precision would continue to increase and decrease, respectively, and the point where they would level off, if at all. Considering that in Experiment 1 RTs did not change with the rotation magnitude – suggesting no mental rotation – we relaxed the control of this variable for Experiment 2. We only made sure that all rotation magnitudes tested in Experiment 1 were repeated at least three times across the experiment and that they added up to 0 to prevent a bias for one side of the rotation. Each sequence length from 2 to 10 was tested fifteen times. In total there were 252 trials divided into: learning to find the center and make reaching movements (48 trials), practice with the target sequences (45 trials; each length tested three times), experiment (135 trials) and washout (24 trials). Apart from these specifications, everything else remained as in Experiment 1.

### Experiment 3 Visuomotor rotation recall (color)

Admittedly, the sequential design of Experiments 1 and 2 departs from what is typically required in a visuomotor rotation task. Therefore, in Experiment 3 (n = 15), we adopted a more conventional trial-pair design to test the working-memory constraints of retrieval strategies. In this design, the participants were first exposed to an observation trial followed by an adaptation trial. On the observation trial, participants were instructed to reach directly to a presented target and to observe the cursor rotation. On the adaptation trial, they were tasked with counteracting the rotation (Figure 1B). These trial-pairs were presented in sequences ranging from 1 to 5 (five targets was the estimated capacity above which guessing began to dominate performance in Experiment 2). After the sequence presentation, there was a memory test in which a target from the sequence was selected and participants had to retrieve the aiming solution for that target to successfully counteract the rotation. In order for participants to easily recognize which type of trial they were in, a text near the center would indicate “Observation”, “Adaptation” or “Memory test” in the corresponding trial.

The target location was fixed at 90° for all trials and target colors were used to indicate the different rotation magnitudes. We chose color to differentiate between rotation magnitudes as it is a feature that has been widely investigated in studies of working memory (Schurgin et al., 2020). For every new sequence, the target colors were sampled without replacement out of 5 possible options: black (rgb = [0,0,0]), orange (rgb = [255,53,51]), green (rgb = [0,255,0]), yellow (rgb = [255,255,0]) and purple (rgb = [178,102,255]). Similarly, the rotation magnitudes associated with each target color were randomly sampled without replacement out of five possible magnitudes: 30, 45 60, 75 and 90°, with randomized signs. Importantly, while participants could experience five possible rotation magnitudes during the sequence of pairs, only three were evaluated in the memory test: ±30, ±60 and ±90. The reason for this was to maintain the duration of the experiment similar to Experiment 1 and 2 while keeping the experiment counterbalanced. Based on previous studies (McDougle and Taylor, 2019) we believed that any effect of the rotation magnitude on performance, e.g. RTs increasing due to mental rotation, would show up with the tested magnitudes. There were a total of 446 trials divided into: learning to find the center and making reaching movements (24 trials), practice with the trial-pair design (16 trials), practice with the sequence of pairs (15 trials), the experiment (375 trials) and washout (16 trials).

### Experiment 4: Visuomotor rotation recall (location)

In order to provide further evidence of the capacity limitations of retrieval strategies in standard visuomotor rotation paradigms, in Experiment 4 (n = 15), we varied the design of Experiment 3 such that instead of colors, each rotation magnitude was associated with a different target location. For every new sequence, the target locations of each pair were randomly sampled without replacement out of eight possible values ranging from 0 to 315° in steps of 45°. Here, the target color was green throughout the experiment. This variation would more closely resemble previous studies of dual adaptation (Baldeo and Henriques, 2013) and pair-trial design (McDougle and Taylor, 2019). Apart from these changes, everything else remained as in Experiment 3, having a total of 446 trials.

### Experiment 5: Visuomotor rotation recall (long-term retrieval)

In this final experiment (n = 15) we aimed to study whether the limited working-memory capacity shown particularly in Experiment 4 (around two elements) would be ameliorated as memories consolidate into long-term storage. In particular, we exposed participants to only sequences of length five and where the target locations were distinct for each rotation magnitude as in Experiment 4. Unlike Experiment 4, this time the rotation magnitude associated with a given target location remained the same across the experiment; however, its position within the sequence was randomized at every new sequence such that participants did not learn a particular arrangement of the trial-pairs. We expected that, over trials, participants would learn the target-rotation associations and guessing behavior would progressively decrease, while memory precision would increase. There were a total of 463 trials, which consisted of: learning to find the center and making reaching movements (24 trials), practice with the trial-pair design (16 trials), practice with the sequence of pairs (22 trials), the experiment (385 trials: 35 sequences) and washout (16 trials). Apart from the above mentioned changes, everything else remained as in Experiment 4.

### Behavioral data analysis

All analyses were performed using the R statistical software version 3.2.2 (R Core team, 2015) or Matlab version 2022a (Natick, MA: The Math Works, Inc., 2022). The core dependent variables in our experiments were reaction times (RTs), target errors, unsigned errors and adapted magnitudes. RTs were defined as the time interval between the target presentation and participants’ departure from the start position. Per-subject medians of reaction times for each condition are displayed in the relevant figures. Target errors were defined as the angular distance (in degrees) between the cursor and the target position. Pooled target errors across subjects are shown in the relevant figures. Unsigned errors refer to the absolute value of the target errors. Per-subject means of unsigned errors for each condition are displayed in the relevant figures. Adapted magnitudes refer to the absolute angular distance of the hand with respect to the target. Per-subject means of unsigned errors for each condition are displayed in the relevant figures. Linear regression models and correlation coefficients were performed using the lm and cor functions on R, respectively. For correlations, we provide both the Pearson and Spearman coefficients. Data points were excluded from analysis if they corresponded to false alarms. A false alarm refers to a trial where participants moved before the target showed up.

### Guessing and memory precision analysis

In order to disentangle the cognitive processes that gave rise to participants’ performance in our tasks, we decided to implement a Bayesian latent-mixture approach (Lee, 2016; Shiffrin 2008). In particular, we assumed that errors in the tasks could be generated by participants sampling from two sources: a memory distribution, represented by a Von Mises distribution, and a guessing distribution, represented by a Uniform distribution – a common approach in studies of spatial working memory (Zhang and Luck 2008). Given that, with the exception of Experiment 1, the number of data points per subject for each sequence length ranged between 3-15, we opted to infer the mixture model parameters in all experiments at the group level to obtain more reliable estimates. The graphical representation of the mixture model for all experiments is shown in Figure 2I. Here, nodes represent the variables in our model and arrows how they influence each other. Shaded nodes are observed variables while unshaded nodes are unknown variables. Circular nodes indicate continuous variables, whereas squared nodes indicate discrete variables. The plates represent independent replications of the graph structure. On the right-side of the graph in Figure 2I we show the model specifications, including prior distributions.

In practice, we implemented the model using the following procedure: for every sequence length, the density from a Von Mises distribution and a Uniform distribution were obtained for every target error. For numerical stability, we used the logarithm of the densities. An indicator random variable *z* for every error and sequence length, selected which of the two distributions was assumed to have generated the target error using their (log) densities and the “ones trick” described in Lee and Wagenmakers (2014). Briefly, the “ones trick” allows us to reliably sample from a target distribution – in our case the mixture distribution – using simpler distributions, given that our target distribution is not available in the inference library.

The indicator random variable *z* was sampled from a Bernoulli distribution with parameter ϕ for every sequence length. This parameter represents the proportion of errors believed to be generated by a Uniform distribution, i.e., the guessing rate. The Von Mises distribution has a mean parameter, µ, and a concentration parameter, κ, for each sequence length. The latter reflects the dispersion in the distribution and in our context represents the memory precision. In order to study serial position effects in our experiments, we varied the structure of this model such that the parameters were estimated per position in the sequence (instead of per sequence length).

In addition to the latent mixture model, we also perform inference on a model that assumes no guessing, i.e., a simple Von Mises distribution. We compared the predictive accuracy of the models using leave-one-out cross validation implemented in R code based on Vethari, Gelman and Gabry (2017).

The posterior distribution of the model parameters was approximated using the software package JAGS (Just Another Gibbs Sampler, Plummer, 2003) implemented in R code. We used three independent chains with 1. 2×10^5^ samples each. A burn-in period (initial samples that were discarded) of 2×10^3^ samples and a thinning of 2 (one every two samples was selected) were used to encourage convergence and reduce autocorrelation between samples. This gave a total of 1. 5×10^4^ posterior samples for each group parameter. The additional JAGS module jags-vonmises (available on https://github.com/yeagle/jags-vonmises) was used to compute Von Mises (log) density values. Convergence of the chains was assessed using the standard potential scale reduction statistic 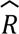 (Gelman and Rubin, 1992). Values of 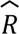 < 1.1 generally indicate that the chains have converged. All the 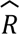 values in our models were below this threshold.

Given that parameter inference was implemented at the group level, and due to the large number of posterior samples, we performed the statistical analyses in the modeling sections using a subsampling procedure (Lehmann and Romano, 2022). In particular, we drew 1000 samples of size b without replacement from the posterior samples of the group parameters, where b was equal to the number of participants for the corresponding experiment. We computed the statistic of interest for each of the 1000 samples and generated a distribution of its value given all samples. We report the 95% confidence interval (CI) based on this distribution. For example, in Experiment 1, in order to test whether there was a correlation between the guessing rate θ and the sequence length, we sampled 15 values of θ without replacement from the posterior distribution of each sequence length (2 to 5 targets), giving a total of 60 data points, over which we performed the correlation test. We performed this step 1000 times and reported the 95%CI of the correlation coefficient based on the resulting distribution.

## Acknowledgments

We thank the members of the IPA Lab for enriching discussions and Justin Jungé for helpful feedback on the statistical analyses. Carlos Velazquez-Vargas and Jordan Taylor were supported by the J. Insley Blair Pyne Fund, Office of Naval Research, Cognitive Science Program and Research Innovation Fund for New Ideas in the Natural Sciences at Princeton University.

## Data availability statement

An abstract of our preliminary findings was presented at the Advances in Motor Learning and Motor Control Proceedings (2023). Data and code from the studies reported in this manuscript are available at https://osf.io/z3cua/

## Supplementary material

**Figure S1:**
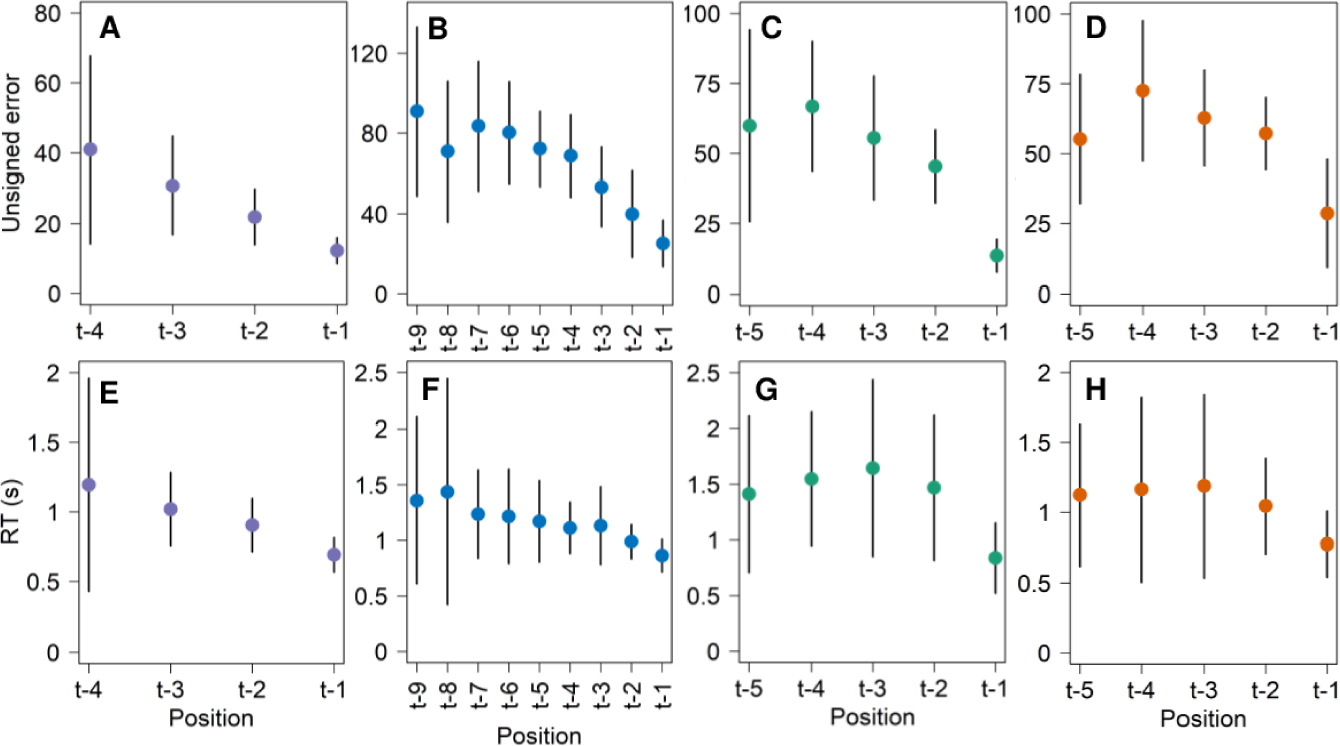
Per-subject serial position effects on unsigned errors (top) and RTs (bottom) in all experiments (purple = Experiment 1; blue = Experiment 2; green = Experiment 3; orange = Experiment 4). Error bars represent the standard deviation with respect to the mean.

**Figure S2:**
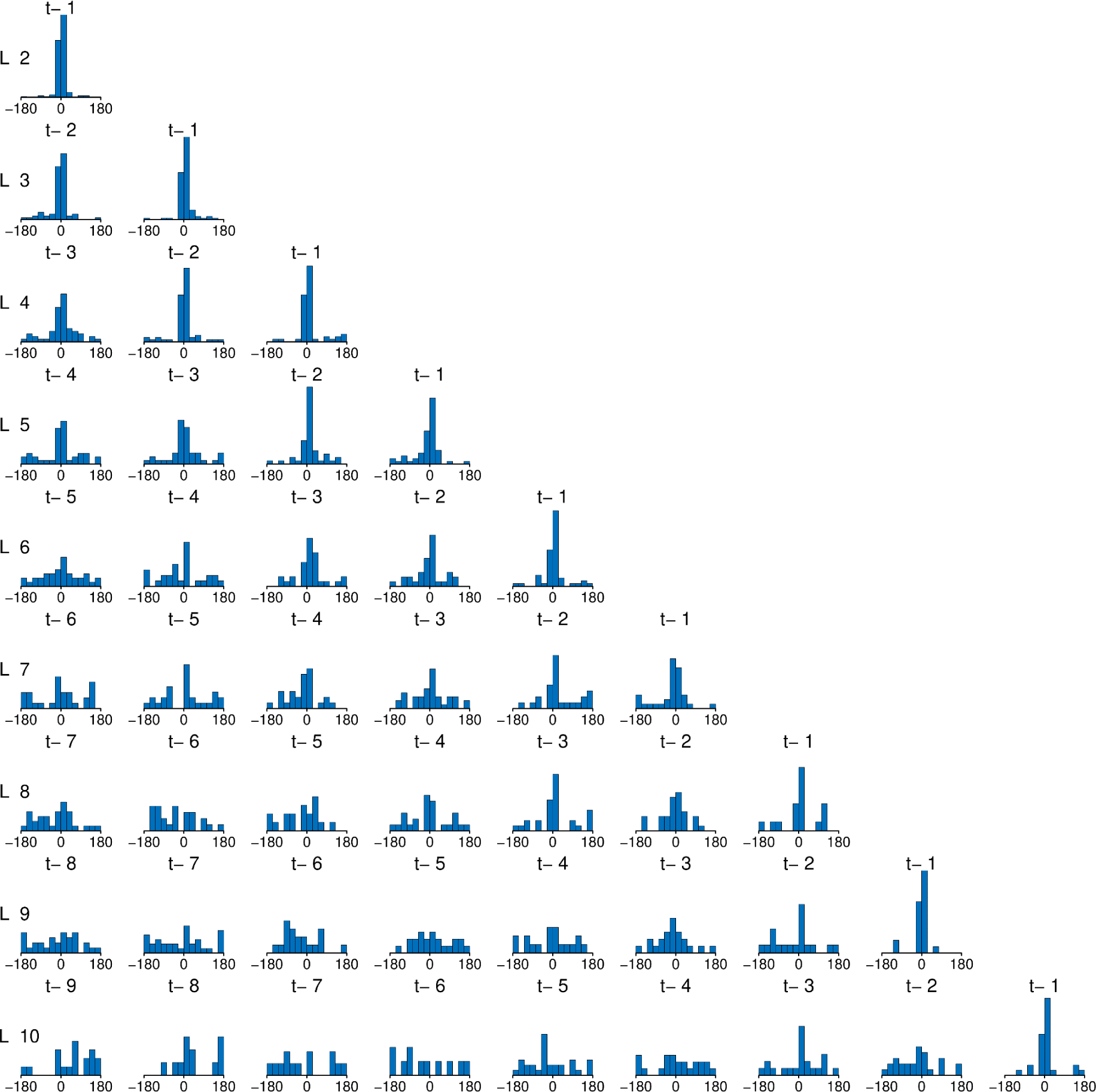
Pooled target error distributions of Experiment 2 by sequence length (rows) and position in the sequence (columns). From right to left, t-1 refers to the most recent item, t-2 to the second most recent item, and so on.

**Figure S3:**
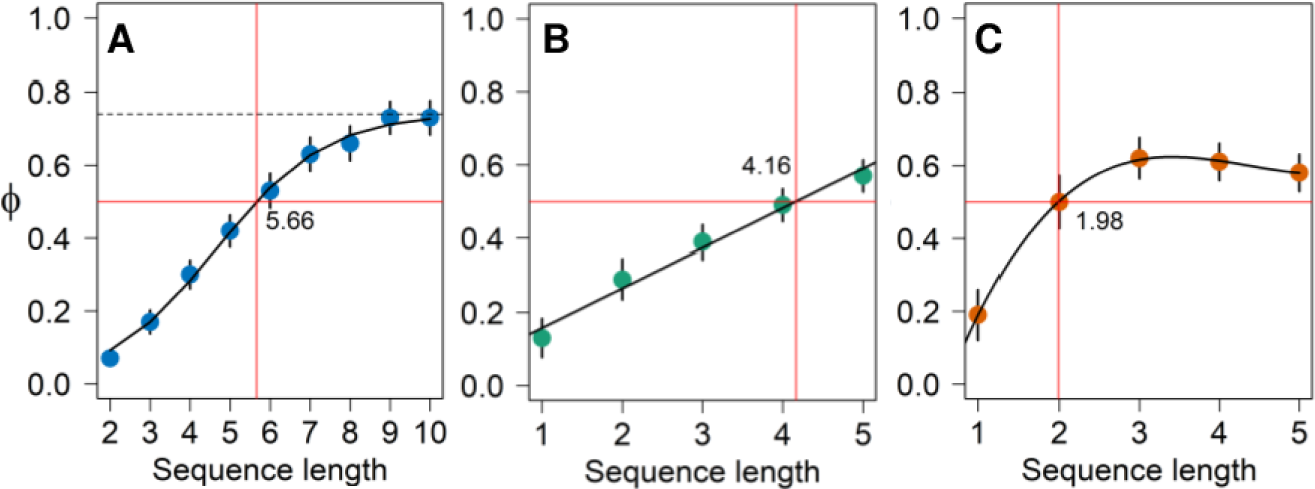
Fit of functions to the posterior means of the group guessing rates θ of Experiment 2 (A), Experiment 3 (B) and Experiment 4 (C) to estimate when θ exceeded 50% of the trials. Red horizontal lines indicate θ = 0.5. Red vertical lines indicate the estimated value of the sequence length where the function crosses this threshold. In panel (A), the black dashed line indicates the asymptote of the sigmoidal function.

**Figure S4:**
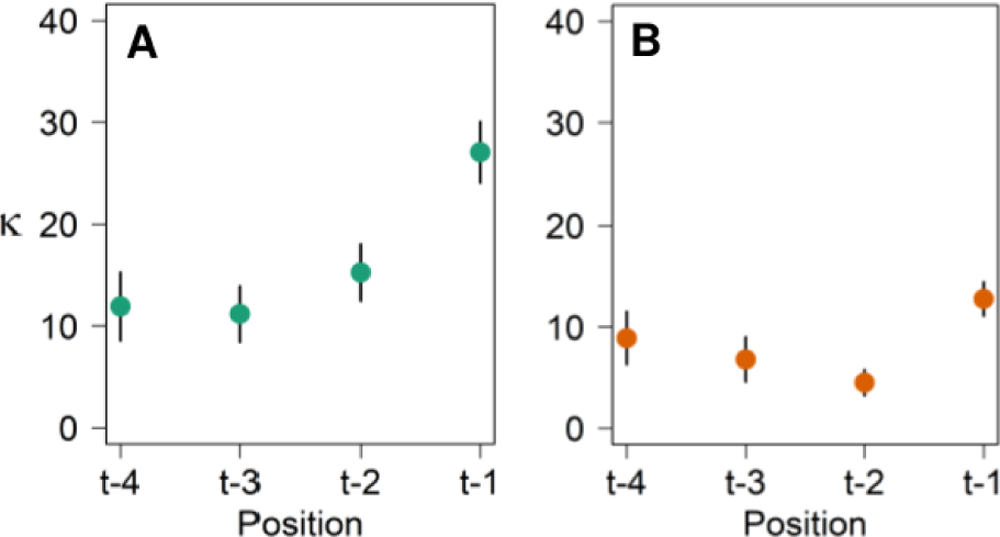
Group serial position effects on memory precision κ in all experiments (green = Experiment 3; orange = Experiment 4). Error bars represent the standard deviation with respect to the mean.

**Figure S5:**
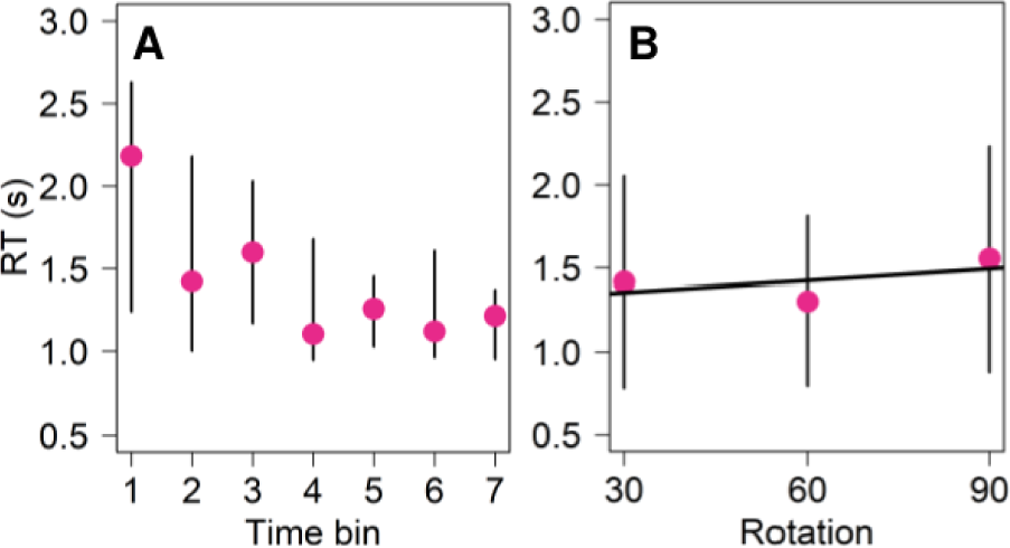
RTs of Experiment 5. **(A)** Per-subject RTs over time bins. **(B)** Per-subject RTs for each rotation magnitude. The black solid line represents the linear fit to the data.

**Figure S6:**
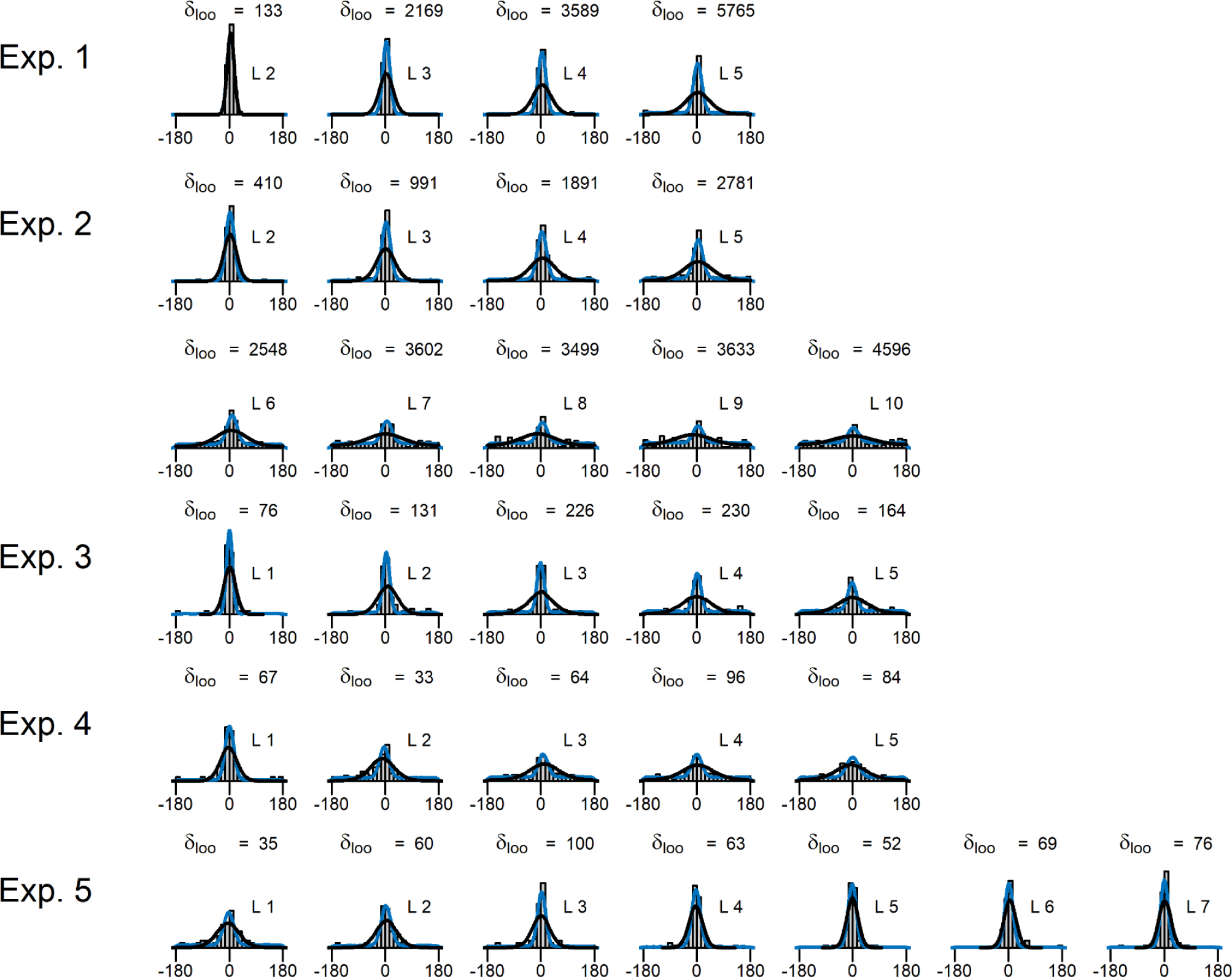
Comparison of the latent-mixture (blue) and the single-source Von Mises (black) models. Densities are generated with the posterior means of the parameters from each model. Histograms represent the pooled target errors. On the top of each panel, we show the difference in the leave-one-one cross validation information criterion (Vethari et al., 2017; Vethari et al., 2023), where positive values indicate evidence in favor of the mixture model.

